# Habitat selection drives dietary specialisation in *Sorex minutus*

**DOI:** 10.1101/2020.02.03.932913

**Authors:** Roselyn L. Ware, Annie L. Booker, Francesca R. Allaby, Robin G. Allaby

**Author notes:** Corresponding author, (RW).

## Abstract

To meet their demand for food, Eurasian pygmy shrews (*Sorex minutus*) require large territories, normally in fields, woodlands, and meadows. Their high metabolism and food requirement often leads to high mortality during winter. However, evidence of shrews in the roof voids of residential buildings has recently been observed, contrary to ecological expectations. Here, five faecal samples collected from different locations were studied by metagenomic analysis to gain information about the shrew’s diets and environments. Two of the samples were collected from novel indoor locations, while the other three were from outdoors in ‘traditional’ habitats. Distinct differences were observed between the diets of the two populations, suggesting a commensal niche expansion has occurred in *S. minutus.* We found that *S. minutus* exploit man-made spaces for foraging, potentially at the cost of a greater parasite burden.

## Introduction

Five species of shrew are known to live in the British isles; the common shrew (*Sorex araneus*), the Eurasian pygmy shrew (*Sorex minutus*), the water shrew (*Neomys fodiens*), the lesser white-toothed shrew (*Crocidura suaveolens*) and the more recently discovered greater white-toothed shrew (*Crocidura russula*) which was found in Ireland in 2007 (1, 2). Here we focus on the Eurasian pygmy shrew (S. *minutus*), which is the smallest British mammal, measuring approximately 8cm from nose to tail and weighing between 2-5g [3]. Despite their small size, shrews have a very high basal metabolic rate [4–6]. However, other factors involving energy usage-such as thermal conductance and body temperature-which are also dependent on body mass [7] have been shown to be as expected given the mass of an average shrew [6].

This high basal metabolism rate requires huge dietary input, with Sorex species having to eat the equivalent of their body mass every day [8]. Shrews, like the rest of the order Eulipotyphla, are insectivores. They are opportunistic foragers and have been recorded to eat a plethora of invertebrates including Isopoda, Chilopoda, Lumbricidae, Diptera, Hymenoptera, Araneae, Mollusca, and more (3, 9). The constant demand for energy means that shrews have adopted a polyphasic circadian rhythm, with around ten active periods per day separated by short rests (10, 11). Activity periods occur during night and day to allow maximum food intake and foraging time. The rest periods rarely, if ever, exceed two hours to prevent starvation [12].

Both the common and the pygmy shrew have large, well defended territories [13]. Although the common shrew (S. *araneus*) is larger than the pygmy shrew (S. *minutus*), their territories are on average smaller. Mean territory sizes have been reported as 1,058 m^2^ for *S. araneus* and 2,146 m^2^ for *S. minutus* [14]. Whilst there are slight differences in habitat preferences between different shrew species in the UK, in general they all require plant cover (provided by smaller plants or trees), access to water and a plentiful supply of food [15]. These criteria are fulfilled by several environments, such as meadows, fields and woodlands. Sorex species (S. *araneus* and *S. minutus*) are generally solitary, however, territorial expansion of females during breeding season allows social contact for reproduction (16, 17). It is likely that this territoriality has developed, in part, as a means of preventing competition for food amongst individuals [13].

Sorex shrews are not thought to hibernate during winter [13, 18, 19]. However, the increasing cold and decreasing supply of prey [3] puts further pressure on individuals to achieve enough energy input to maintain their high basal metabolism. Common adaptive strategies seen in other small mammals, such as huddling [20], cannot be adopted by Sorex shrews due to their territoriality, particularly in winter when little to no contact is observed between individuals [21]. Several adaptations have developed in order to survive during the winter, both behavioural and physiological [22]. Nests are used throughout the year, allowing a reduction in thermogenic cost whenever the ambient temperature is lower than approximately 24 °C, above which point the effects are negligible as thermoneutrality is reached [18].

Over the past 3 years we have retrieved evidence of Sorex species in residential building roof void spaces whilst sampling for bat guano. Whilst previous observations have been documented of urban occurrences of shrew (23, 24) this is not seen as common behaviour and far removed from the common, natural habitats of woodland or grassland [15]. Both *S. minutus* and *S. araneus* require large territories of between 1-2 km^2^ [14] and *S. araneus* requires ground in which it can burrow to search for prey (9, 25). Neither of these requirements are fulfilled by the environments in which evidence of activity has been found. It is therefore possible that Sorex shrews may be seeking temporary shelter as a method of avoiding harsh climates; avoidance tactics have previously been recognised during glaciation events through the Pleistocene era (26, 27). In this study, the environments and diets of shrews were analysed through metagenomic analysis of faecal samples found in a range of locations, with the aim to determine any differences between samples found at conventional and novel sites to gain insight into whether the urban environment represents a dietary niche or just shelter from unfavourable conditions. Emphasis was placed on determining any differences between the diets of shrews from the different environments, with an overall aim of determining whether the shrews which were found indoors were hunting in their novel environments, or whether they were only temporarily sheltering indoors and predominantly hunting outside.

Sorex shrew diets have never been previously been determined using a metagenomic approach, with most of the extant data coming from older studies, before the technology was available [8, 9, 28–31]. Previous study techniques generally involve analysis of excised digestive tracts or faecal samples by microscope [8, 9, 30]. This only allows analysis of a very small portion of the diet that has not been fully digested, as the sample must be intact enough to be recognised [32]. A metagenomic study allows unbiased, broad spectrum, non-invasive study of individuals in their natural environment (33, 34). It also allows study of the environment through types of environmental contamination introduced at the sample site [35]. For example, particular fungal, bacterial or plant matter present may provide more information of the location or behaviour of the shrew. Metagenomic studies also allow relative quantification, which may elude to the proportion of particular types of prey in the diet of individuals [34].

## Methods

In this study, five faecal samples determined to be from *S. minutus*, were studied. The samples each came from different locations, detailed in Table S1 (supplementary material). All faecal samples were submitted to the University of Warwick as part of the EcoWarwicker Ecological Forensics service, with those collecting the guano responsible for obtaining all necessary permits. All the work described here was approved by the University Genetic Modification and Biosafety Committee and the ethical issues were approved by the University Animal Welfare & Ethical Review Body Committee. Anonymised samples were used in this project with the consent of EcoWarwicker Ecological Forensics. As faeces was the source of the material, no animals were handled or disturbed in the completion of this project.

DNA was extracted as follows. Samples were crushed, then incubated in CTAB buffer at 37°C overnight. The incubated samples were then centrifuged for 1 minute at 25,000 g, and the supernatant retrieved. 300 μl of chloroform:isoamyl alcohol (24:1) was added to each sample before vortexing, centrifuging 1 minute at 25,000 g, and retrieving the supernatant. DNA was purified using the Qiagen MinElute PCR purification kit, and then quantified using the Qubit™ dsDNA HS kit.

The species was confirmed through PCR and sanger sequencing; 20 μl PCRs were prepared using the primers shown in Table S2, each at 5 μM. The reactions contained 2 μl 10X Platinum^®^ Taq buffer, 2 μl of dNTPs at 2mM, 0.8 μl 50mM Mg^2+^, 1.3 μl primer mix, 0.1 μl Platinum^®^ Taq DNA polymerase, between 0.2-2 μl of sample (dependent on concentration of extract) and 11.8-13.6 μl ultrapure H_2_O. Touchdown thermal cycling conditions were as follows: 5 mins at 95°C, followed by 10 cycles of 94°C for 30s, 57°C for 30s (decreasing by 0.1°C per cycle) and 72°C for 30s, followed by 32 cycles of 95°C for 30s, 54°C for 30s, then 72°C for 30s, followed by a final extension period of 72°C for 7 minutes. Reaction success was confirmed by electrophoresis on a 2% agarose gel, clean-up was undertaken by adding 2 μl of Fast-AP and 0.5 μl of Exonuclease-1, then incubating at 37°C for 30 mins, then 80°C for 15 mins. Forward primers (Table S2.a) were used in a GATC LightRun™ Sanger sequencing reaction. The sequences (Table S3) produced were BLAST searched against the nucleotide database to determine the species [36].

Library preparation was adapted from the protocol described in Meyer et al. [37], with the following adaptations described in [38]. 0.1 μl of adapter mix used during adapter ligation, spin column purification instead of SPRI (MinElute PCR purification, Qiagen), the purification step after adapter fill-in replaced with heat inactivation for 20 minutes at 80 °C, and the use of dual indexing. The protocol was also modified as follows: No fragmentation step, the volume of the Blunt-end repair reaction was 40 μl, the T4 DNA ligase was added to individual sample tubes instead of the master mix during adapter ligation, Platinum Pfx was used for indexing PCR. Index sequences were selected such that a there was a minimum Hamming 12 distance of 3 between any given two of the forwards (5’) and the reverse (3’) indexes. The indexes selected are shown in Table S4.

All stages of preparation before the indexing PCR were carried out in a dedicated DNA laboratory that deals specifically with non-amplified samples to prevent contamination. A 2% agarose gel was then set up and run to confirm the presence of DNA in each of the libraries. The libraries purified using solid phase reversible immobilisation (SPRI) beads [39].

Libraries were quantified using the Qubit™ dsDNA HS kit, then characterised using the Agilent High Sensitivity Bioanalyzer in order to calculate the molarity. Libraries were then normalised to 4nM, and pooled. The pool was spiked with 1% PhiX control v3 library. The pooled libraries were sequenced using the Illumina MiSeq using a 300 cycle V2 reagent cartridge [40].

Demultiplexing was carried out by the Illumina MiSeq system. Initial data was first visualised using FastQC (version 0.11.6). Adapter sequences were removed and the forward and reverse sequences were collapsed using AdapterRemoval (version 2.2.2) [41], specifying a minimum length of 30 and a minimum quality of 30. The fastq files were then converted to fasta files using awk. Duplicates were assumed to be an artefact of PCR and were removed using the fastx_collapser command from the FASTX-toolkit (version 0.0.13) [42]. A metagenomic BLASTn search (version 2.6.0) was undertaken to confirm species identification (Zhang, 2000). In order to remove falsely assigned reads, reads that had been assigned to bacteria, Mammalia, fungi, Protostomia, and Viridiplantae were subjected to Phylogenetic Intersection Analysis (PIA) as described by [43]. Taxa with >2% of the found reads also occurring in the blank were removed.

## Results and discussion

Samples A-E were confirmed as droppings from *Sorex minutus* by PCR and sequencing of a portion of the MT-CYB gene (Table S3). An unbiased shotgun approach was used to sequence DNA from the droppings to identify dietary components on the MiSeq platform. Of a total of 14.9 million reads returned, 832,223 reads had positive assignations to Protostomia, Mammalia, Viridiplantae, fungi, and bacteria before PIA (Fig 1A), with 166,328 after PIA (Fig 1B). PIA significantly reduces the number of assignations (S5 and S6 Table), but largely does not alter the broad profiles of assignations for each sample.

**Figure 1.**
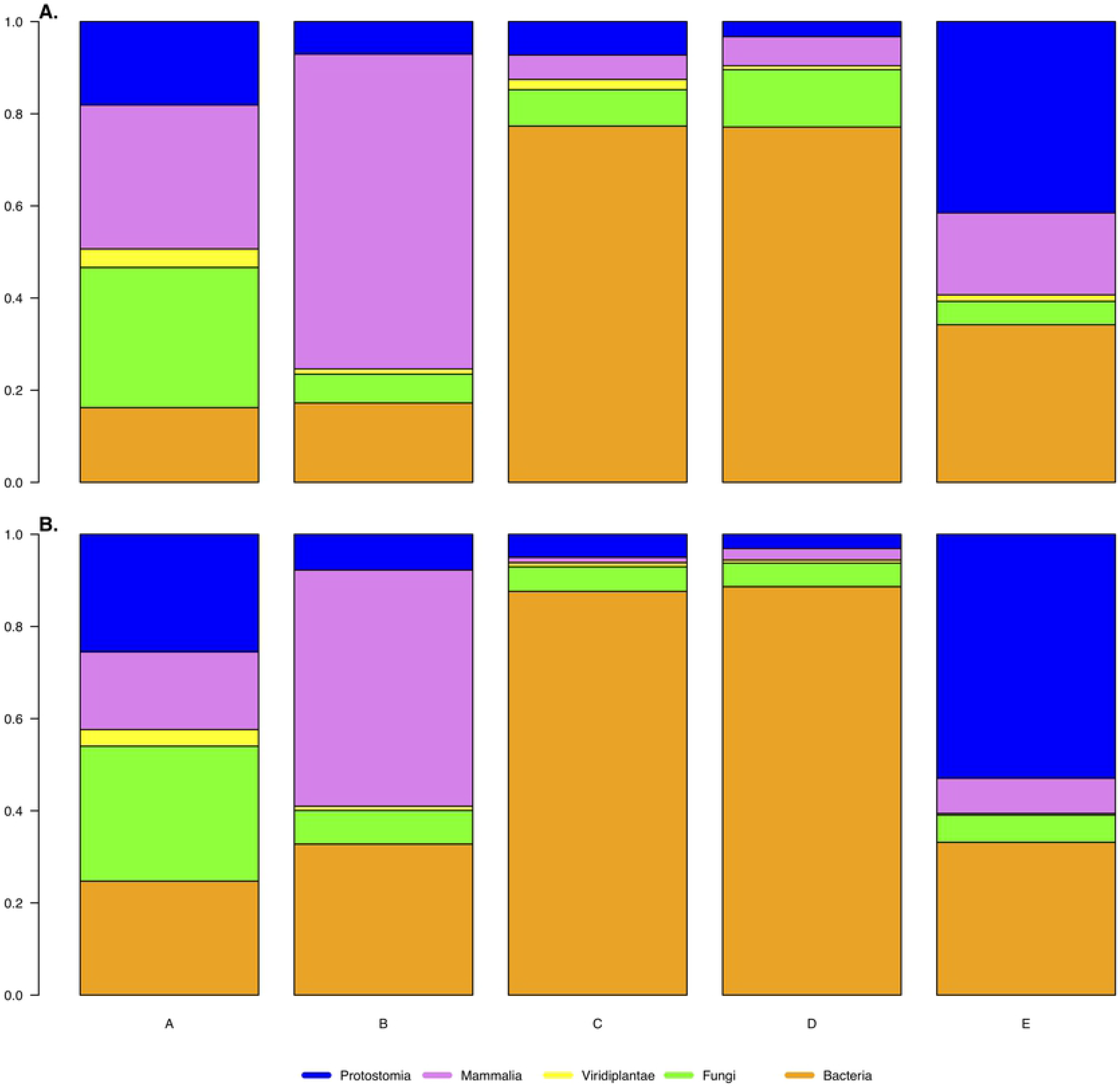
The proportions of reads assigned to Protostomia, Mammalia, Viridiplantae, Fungi and Bacteria. (A.) before and (B.) after PIA. Samples were collected from locations that were indoor (samples A and B) or outdoor (samples C, D, E).

Bacteria is the most abundant of the major groups across all samples. The presence of bacteria is due to several factors, namely the gut microbiota of each individual, and environmental contamination from the site at which the sample was found.

The presence of mammalian reads is largely due to the individual that produced the original faecal sample (Fig 1). The most predominant read in the group Mammalia, across all samples except the blank, was Insectivora of the family Soricidae. The metagenomic approach was able to identify the likely source of the samples as Sorex in the case of all samples, rather than *S. minutus*. This is because the whole genome data of *S. minutus* is not currently available. Initially, the DNA was matched to the closest available genome, which happens to be *S. araneus*, however, the PIA was able to assign 1,730 reads to Sorex (the intersect between *S. araneus* and *S. minutus*).

Another major group detected is fungi. The presence of fungi in samples is assumed to be almost entirely from the environment. The presence of Viridiplantae is also expected to be predominantly as a result of the environment, as *S. minutus* is an insectivore.

Protostomia present in the sample is assumed to largely be due to dietary input (Fig 2 However, the clade also includes nematodes and other non-dietary taxa. The presence of these groups is expected to be partially due to environmental contamination, and also due to some potentially parasitic species. For example, the order Strongylida is known to contain many parasitic and pathogenic species [44], while the order Rhabditida contains many species which live in soil [45].

**Figure 2.**
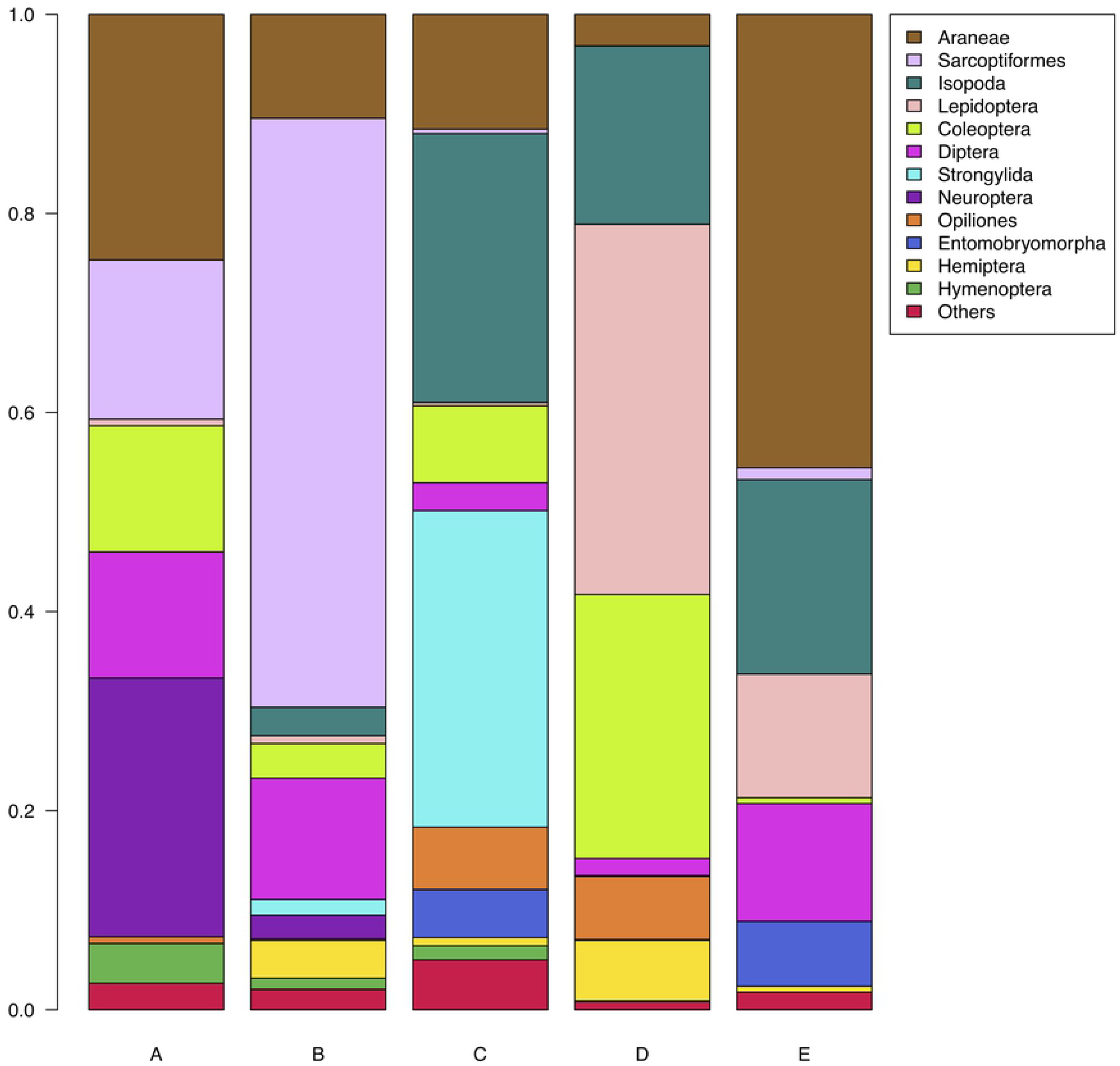
The phyletic distribution of reads assigned to Protostomia at order level after PIA. Taxa comprising <1% were collapsed into “others”. Samples were collected from locations that were indoor (samples A and B) or outdoor (samples C, D, E).

Fig 2 shows the phyletic distribution of reads assigned to Protostomia (the likely prey of shrews) collapsed to order level. Previous studies have suggested that *S. minutus* has a broad diet, largely comprised of Araneae, Opiliones, Coleoptera, and Isopoda (47, 49), all of which are observed in this study, although Opiliones comprised a smaller proportion of the diets in this study. As has been previously observed (47, 49), Annelids (earthworms) did not form a large portion of the diet, possibly due to the small body size of *S. minutus*. Additionally, significant proportions of the diets here were comprised of Lepidoptera, Diptera and Neuroptera.

The main prey of the loft sample, sample A, appears to be of the class Arachnida (Araneae (spiders) and Sarcoptiformes (mites)), followed by Neuroptera (net-winged insects). Significant amounts of Diptera (flies), Coleoptera (beetles), were also detected. Spiders being common in houses, this suggests they may have been hunted within the building. Similar to sample A, the main dietary components in sample B, collected from a barn, was also of the class Arachnida but with a more substantial Sarcoptiform component, with signals also present for Diptera. Another substantial dietary component that was detected was Sarcoptiformes, an order containing many species of mites. The sample found in a barn (number B) also contains a large proportion of data assigned to Sarcoptiformes, whereas the other samples (collected from outdoor locations) had little or no sarcoptiform DNA. It may be that transmission of mites occurs more readily in indoor environments.

Fig 3 shows the phyletic distribution of reads assigned to Viridiplantae collapsed to order level (panel A) and Genus level (panel B). Sample C was found in the bark of a tree. From the DNA data it is likely that it was found in a tree belonging to the genus Quercus (oak) (fig 3. B). The same sample contains a very high abundance of bacteria belonging to the order Pseudomonadales, specifically to the *Pseudomonas fluorescens* group, which are typically plant associated. Sample D was found in a campsite, set in meadowlands with several lakes nearby. Whilst it is known that British shrews can swim [46], it is not clear whether shrews will seek out aquatic prey by hunting in water: whilst *Neomys* shrews are known to forage in water, it has not been observed in *Sorex* [47]. It is possible that the individual from which sample D came from, may live close to one of the lakes on the site as this sample was shown to contain significantly more algae than each of the other samples. It is possible that this environmental contamination due to being found close to a body of water, but there is a possibility that it may be present due to consuming prey which had consumed, or been covered in algae.

**Figure 3.**
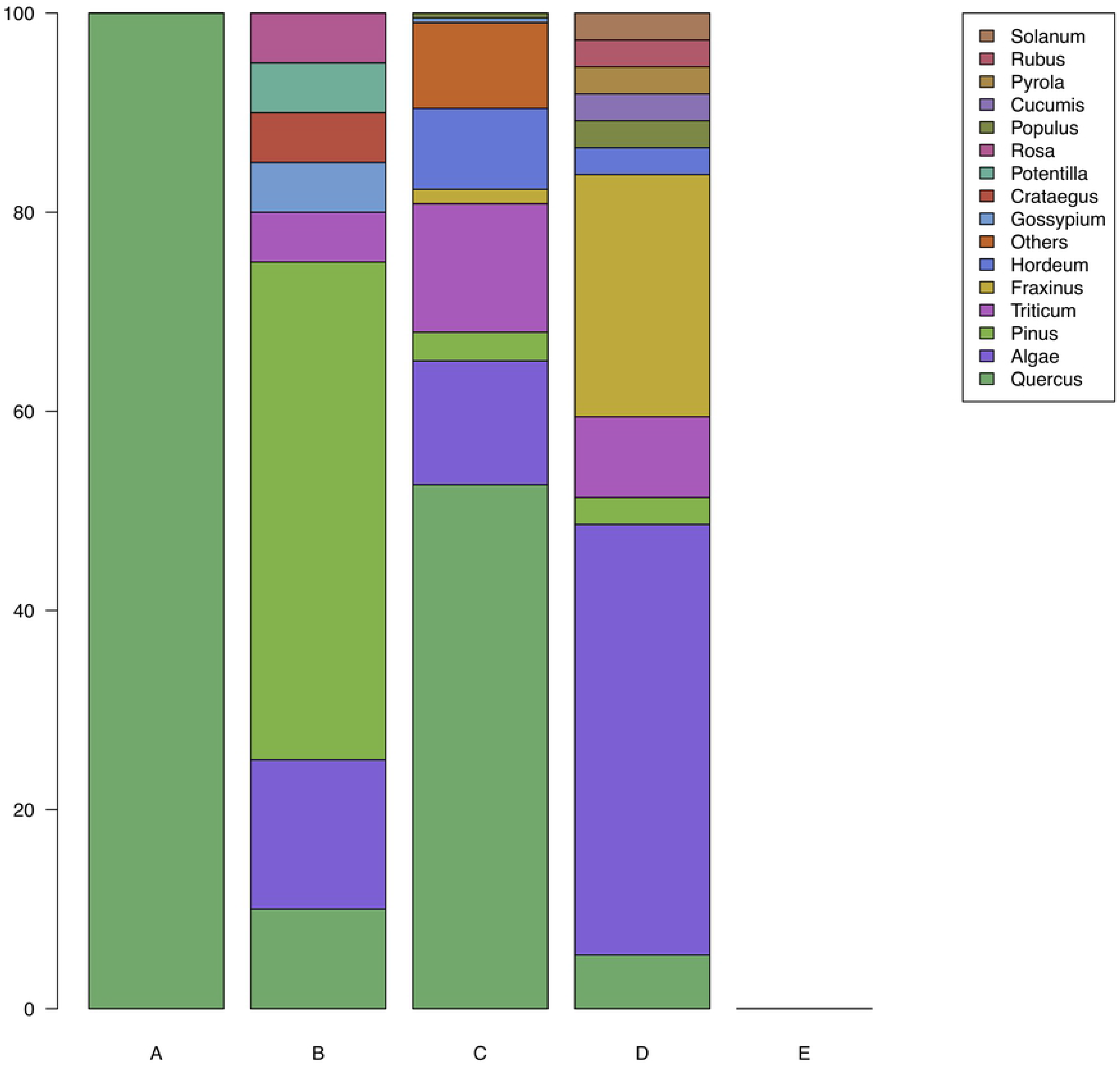
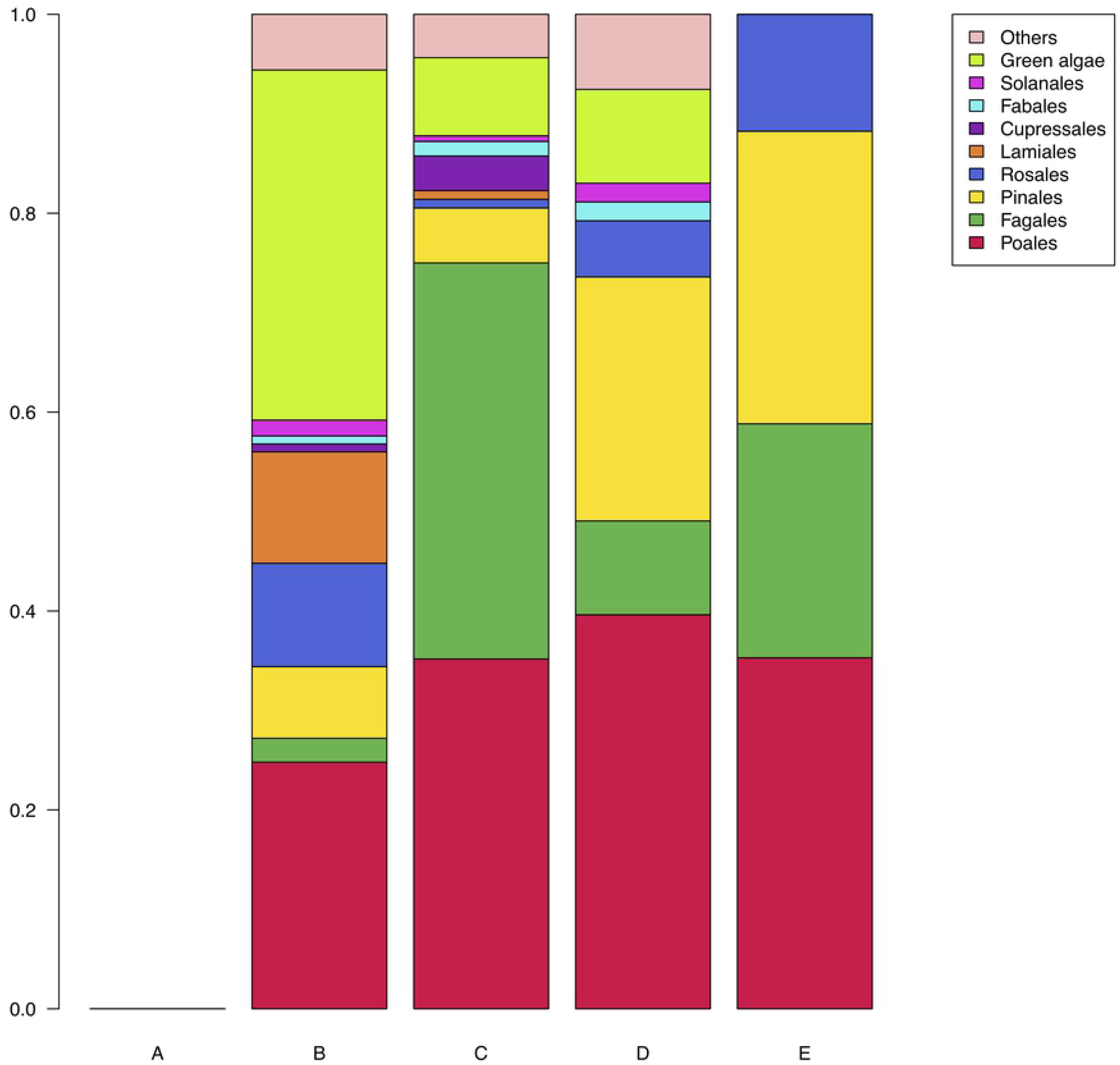
The phyletic distribution of reads assigned to Viridiplantae at (A) order level, and (B) Genus, after PIA. Taxa that appeared in the blank at >=2% of the assigned reads were removed. Taxa comprising <1% were collapsed into “others”. Samples were collected from locations that were indoor (samples A and B) or outdoor (samples C, D, E).

Sample number C, which was discovered in the bark of a tree, shows a large proportion of Isopoda (woodlice), followed by Araneae, meaning that the diet likely composed of mostly woodlice and spiders, both common inhabitants of woodland environments, supporting the data that suggests that *S. minutus* forages terrestrially, even when in close proximity to water [47]. Whilst there is a large number of reads assigned to Strongylida (nematodes), this is unlikely to be a dietary component. Isopoda also formed a substantial proportion of the other two outdoor samples (D and E).

The Protostomia data returned for sample D, found in a campsite with abundant meadowlands and lakes, is dominated by reads assigned to Lepidoptera (moths and butterflies). Considerable numbers of reads were also assigned to Coleoptera and Isopoda. The order of highest abundance detected in sample E, found outside a church in a small village, was Araneae, with Lepidoptera, Isopoda, and Diptera.

The relative diversities (Shannon-Weaver), shown in fig 5, suggests that the most diverse sample is sample C (from inside the loose bark of a tree), followed by sample A (from Loft void in a residential building). The least diverse is sample B (from inside a barn). A comparison of the niche breadth of all reads in each sample are shown in the plot in fig 5. The Levin’s index plot largely confirms the Shannon-Weaver plot in fig 4. However, the difference in diversity between each sample is more pronounced when looking at the Levin’s index plot as opposed to the Shannon-Weaver plot.

**Figure 4.**
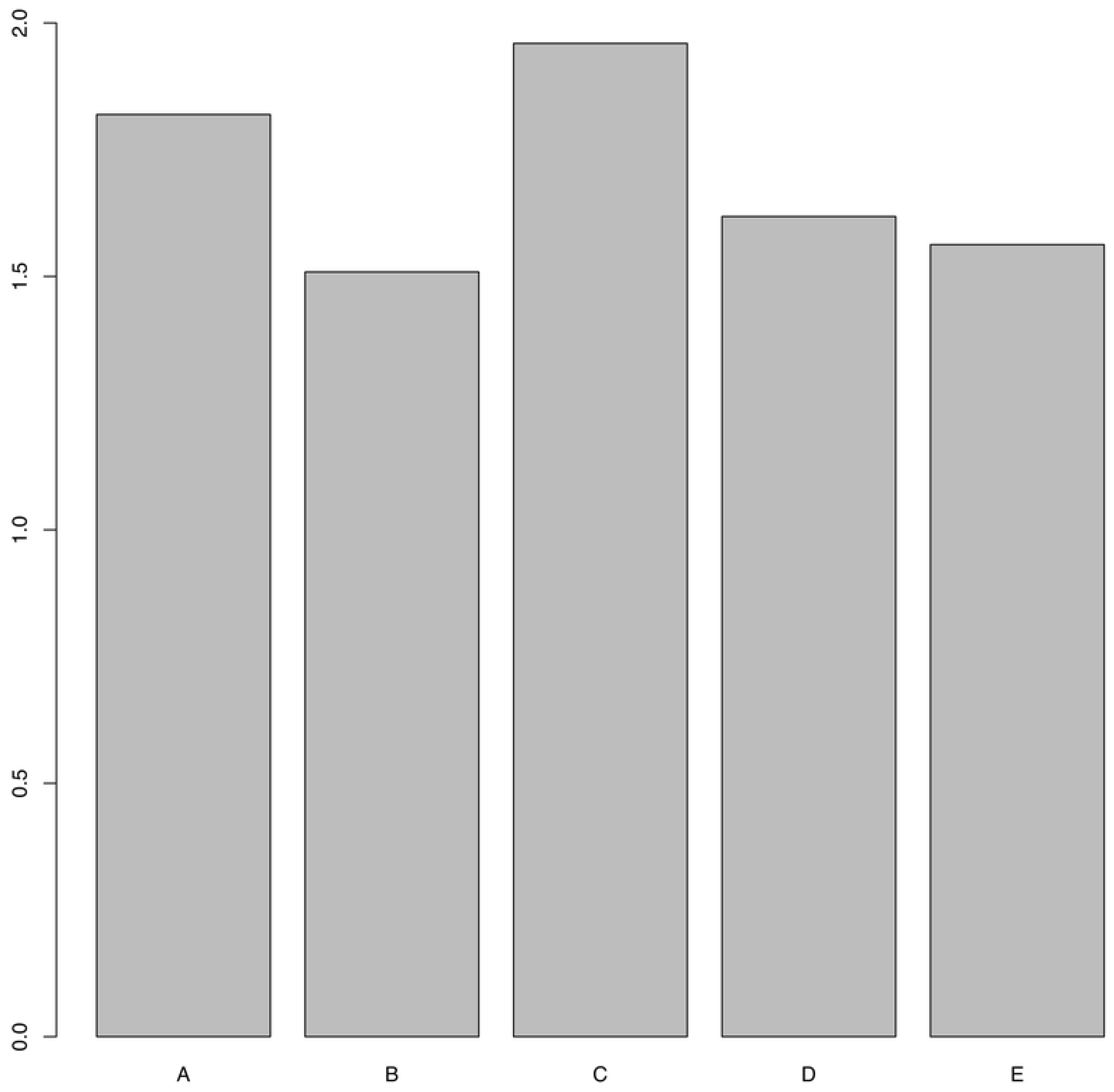
Shannon-Weaver index values for Protostomia collapsed to order level, for each sample, after PIA. Samples were collected from locations that were indoor (samples A and B) or outdoor (samples C, D, E).

**Figure 5.**
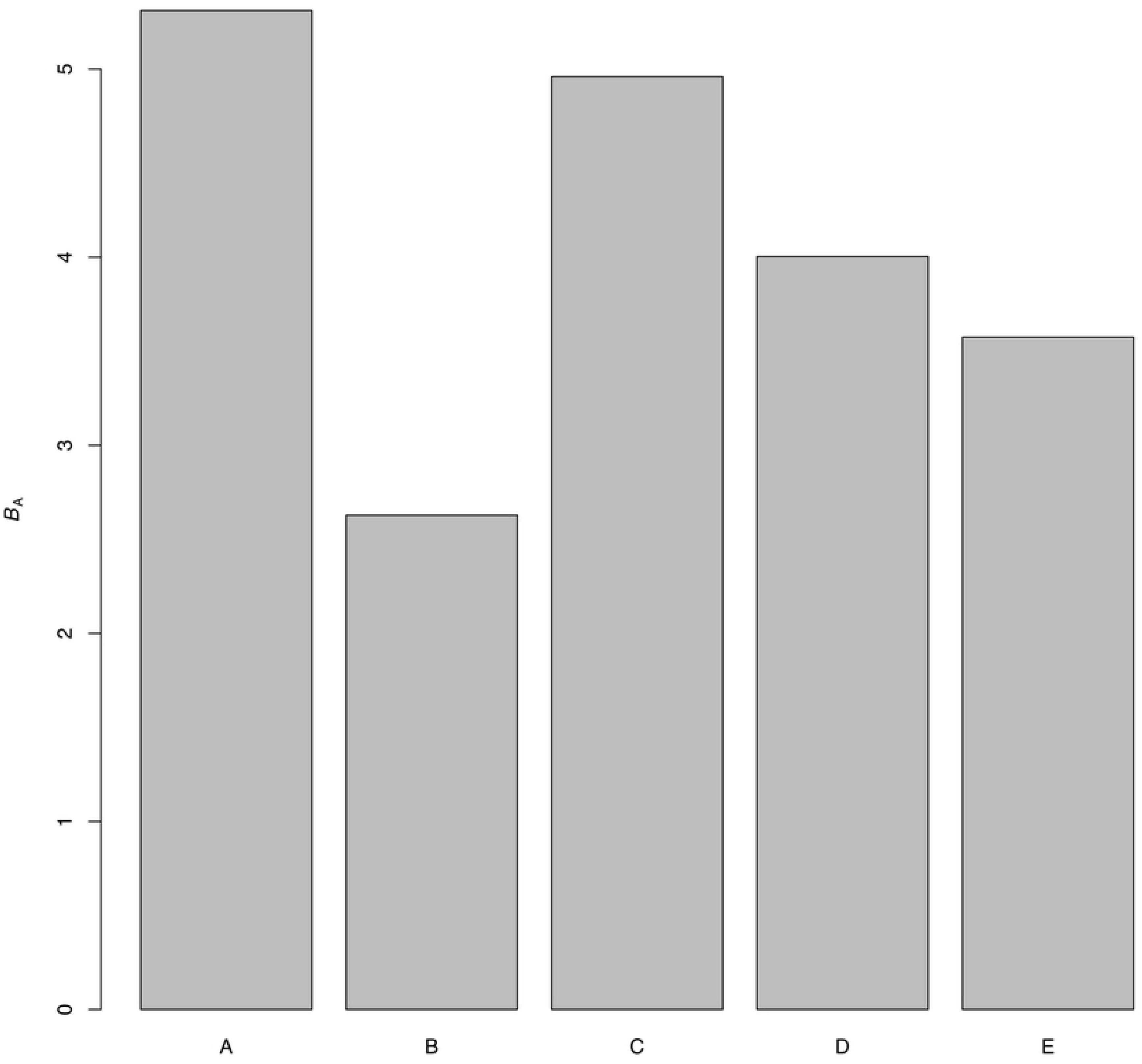
A comparison of the Levin’s index values calculated for each sample, after PIA. The index value, also known as relative niche breadth, is equivalent to relative diversity. Levin’s indexes account for differences in abundance between different taxa. Samples were collected from locations that were indoor (samples A and B) or outdoor (samples C, D, E).

Pianka’s index values for the total assigned reads is shown in fig 6 indicating the extent of dietary overlap between samples. This analysis suggests two dietary guilds or niches which correlate with commensal and natural environments respectively. In the first, samples A and B (collected inside a roof void and barn respectively) correlate closely. The second guild appears to be broader encompassing the outdoor samples, which show some degree of niche overlap.

**Figure 6.**
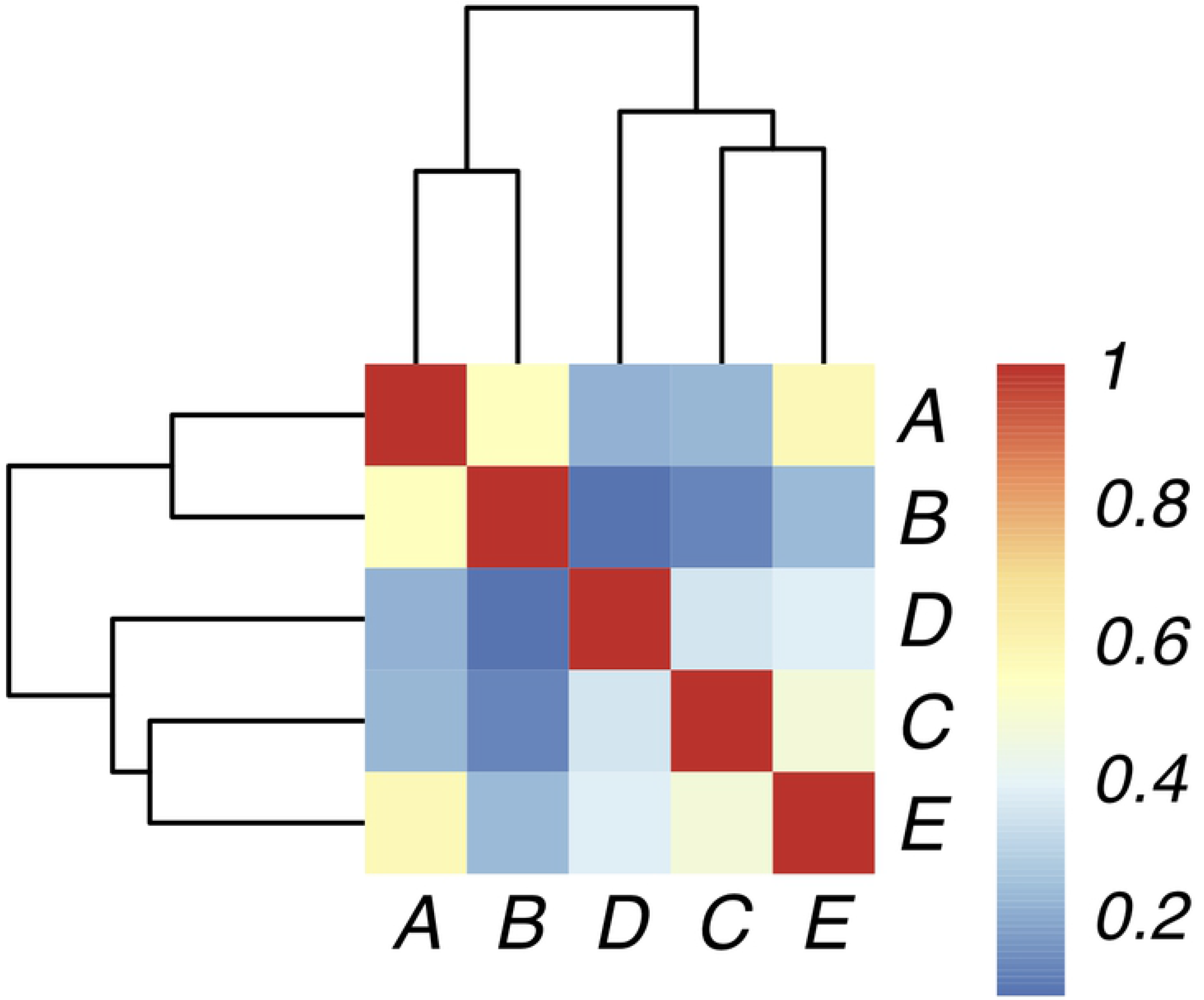
Pianka’s overlap indexes comparing the total overlap of all data obtained from each sample at order level, after PIA. A value of 1 means two samples overlap completely and are therefore identical, whereas a value of 0 means there is no overlap between two samples and they are completely distinct from one another. Data sorted using complete-linkage clustering using the R hclust package [48]

## Conclusions

Whilst faecal metagenomics has been previously done on other species either to obtain information about gut microbiomes [50] or to learn more about individuals’ diets (32, 34), this metagenomic approach to diet profiling has never before been carried out on Sorex shrews. This project has allowed the generation of a vast amount of very informative data, both about the diets and the environments of five individual British pygmy *Sorex minutus*. The aim of the project was to determine whether there were differences between the diets of shrews for which faecal samples were found in ‘traditional’ habitats (i.e. in meadows, woodlands or grassy gardens) and in ‘novel’ habitats (i.e. barns and residential buildings). The purpose of this was to determine whether shrews which are being discovered indoors are sheltering temporarily or are living in such locations more permanently. The data which was obtained gave a highly useful indication as to the diet of each individual. While each diet was unique, with different types of prey forming different proportions of the diets, there were more similarities between the diets of the indoor shrews than with the outdoor shrews and vice versa. This distinction between the diets of the indoor and outdoor shrews suggests that those individuals depositing faeces inside may also be living and foraging more permanently inside, and that the habitat selection has driven intraspecies dietary specialisation within *S. minutus*. We also see that habitat selection may impact parasite burden, with indoor individuals bearing a much greater proportion of parasite load.

## Data availability statement

The datasets analysed for this study can be found in the European Nucleotide Archive under the project code PRJEB35343. See supplementary Table S7 for sample accession codes.

## Author contributions

Study design by RA, RW, FA, and AB; lab work undertaken by AB with supervision by RW and FA; data analysis by RW and AB; manuscript written by AB and RW with review and editing by RA.

## Supporting information

**S1 Table. The identification numbers given to each sample in the study and the location in which each sample was collected**.

**S2 Table. Barcoding PCR primers**

**S3 Table. Confirmation of *Sorex minutus* ID using sanger sequencing**

**S4 Table. Illumina I5 and I7 indexes**

**S5 Table. Taxa counts pre-PIA and post-PIA**

**S6 Table. Read counts pre-PIA and post-PIA**

